# Hidden in plain sight: What remains to be discovered in the eukaryotic proteome?

**DOI:** 10.1101/469569

**Authors:** Valerie Wood, Antonia Lock, Midori A. Harris, Kim Rutherford, Jürg Bähler, Stephen G. Oliver

## Abstract

The first decade of genome sequencing stimulated an explosion in the characterization of unknown proteins. More recently, the pace of functional discovery has slowed, leaving around 20% of the proteins even in well-studied model organisms without informative descriptions of their biological roles. Remarkably, many uncharacterized proteins are conserved from yeasts to human, suggesting that they contribute to fundamental biological processes. To fully understand biological systems in health and disease, we need to account for every part of the system. Unstudied proteins thus represent a collective blind spot that limits the progress of both basic and applied biosciences.

We use a simple yet powerful metric based on Gene Ontology (GO) biological process terms to define characterized and uncharacterized proteins for human, budding yeast, and fission yeast. We then identify a set of conserved but unstudied proteins in *S. pombe*, and classify them based on a combination of orthogonal attributes determined by large-scale experimental and comparative methods. Finally, we explore possible reasons why these proteins remain neglected, and propose courses of action to raise their profile and thereby reap the benefits of completing the catalog of proteins’ biological roles.

## Slow progress in characterizing unknowns

When the first eukaryotic chromosome (chromosome III of *Saccharomyces cerevisiae*) was sequenced in 1992, the most surprising discovery was that previously undetected protein-coding genes outnumbered mapped genes by a factor of five [1, 2]. Researchers had generally assumed that few proteins remained to be discovered, especially in an organism as intensively studied as yeast. The completion of the *S. cerevisiae* genome sequence in 1996 confirmed that more than half the genes lacked any indication of their biochemical activity or broader biological role [2]. Over the ensuing two decades, complete genome sequences have become available for over 6500 eukaryotic species [3]. At first, characterization of newly discovered genes progressed rapidly in model species, as researchers supplemented classical biochemistry and forward genetics with reverse genetics, homology modeling, and large-scale systematic techniques to study novel genes. Complete genome sequences also allowed the deployment of large-scale systematic techniques [4]. More recently, however, progress has slowed, even in well-studied species such as the budding yeast *S. cerevisiae* and the fission yeast *Schizosaccharomyces pombe*.

Fig 1 shows protein characterization over the past 28 years for these two yeasts. Notably, the proportions characterized in fission yeast (84%) and budding yeast (82%) have only slightly increased in the past decade (from 80%, as noted by Peña-Castillo and Hughes [5], and 77% respectively). Across all studied eukaryotic species, the proportion of characterized proteins has reached a plateau around 80%, and exhibits a long-tailed distribution, with the biological roles of the remaining 20% still elusive.

**Figure 1:**
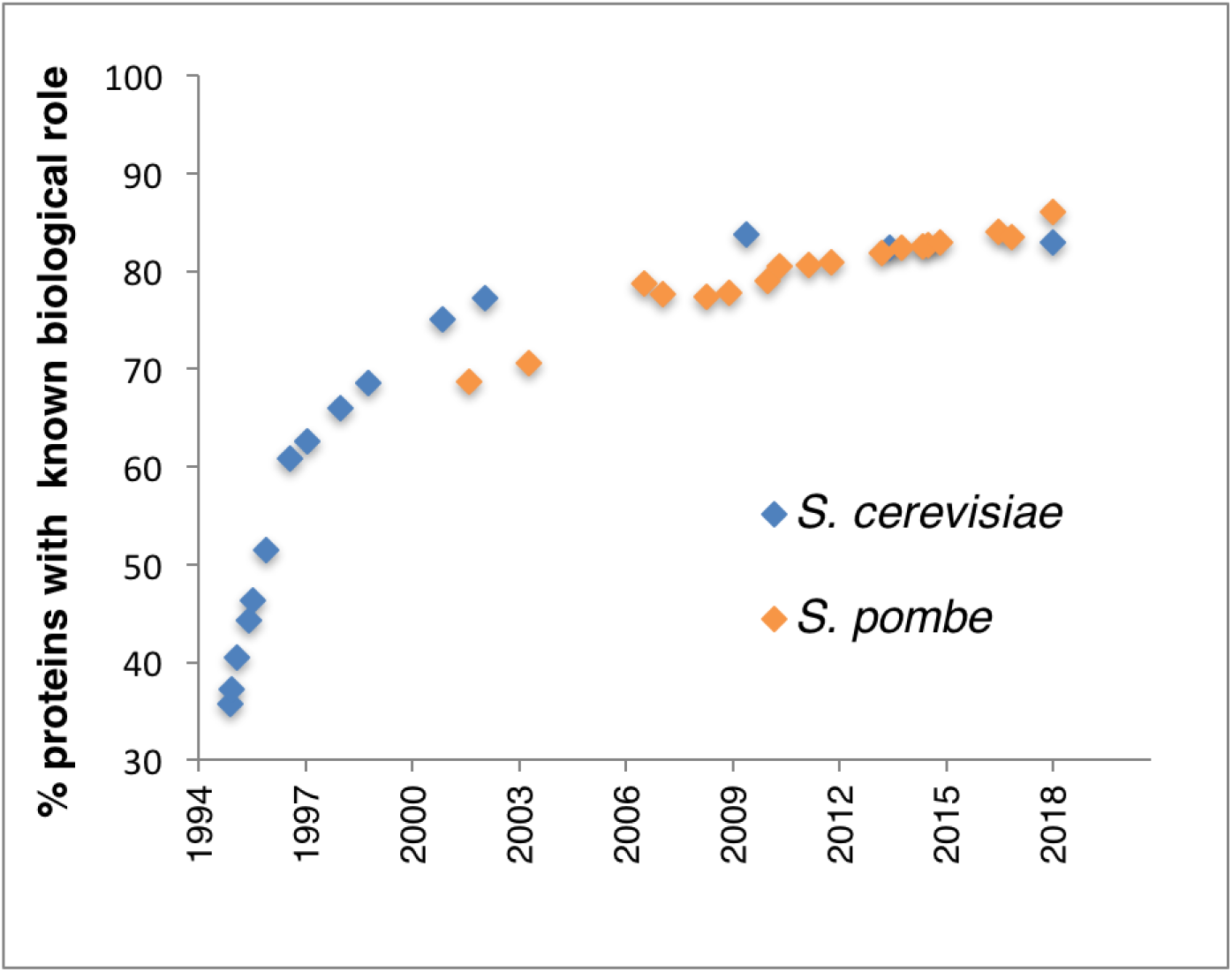
Characterization history of budding yeast and fission yeast proteins. Numbers of *S. pombe* and *S. cerevisiae* proteins that have had their biological roles either determined from experiments or inferred from sequence orthology to known proteins in other species, plotted as a function of time. The numbers of unknown proteins have not markedly decreased over the past 15 years. Data sources: *S. cerevisiae* 1994–1998 [22], 2000 [23]), 2002[24], 2009 [25], 2013 [26], 2018 this study (see Figure 3); *S. pombe* [27].

Here we use a simple yet powerful metric based on Gene Ontology (GO) biological process terms to define characterized and uncharacterized proteins for human and the two model yeasts. We then combine our GO-based classification with information about taxonomic conservation using fission yeast to identify a set of broadly conserved, but unstudied, proteins. We classify the fission yeast conserved but unstudied protein set based on a combination of orthogonal attributes (e.g. taxonomic conservation, mutant viability, protein sequence features, localization). Finally, we explore possible reasons why these proteins remain neglected, propose courses of action to raise their profile among bench researchers and bioinformaticians, and posit the benefits of completing catalog of proteins’ biological roles.

## Defining unknown metrics: what counts as “known”?

To estimate more precisely the proportion of a proteome that is characterized, and to provide inventories of uncharacterized gene products, the “known” category must be rigorously defined. However, the gradual accumulation of data of many different types, from diverse experimental and computational methods and multiple sources, makes it challenging to draw a clear line between “known” and “unknown”. For example, in 2004 Hughes *et al*. [6] observed that the then-current Yeast Proteome Database (YPD) listed 80% of *S. cerevisiae* genes as “known”, but also noted that by more stringent criteria based on Gene Ontology (GO) annotation then in the *Saccharomyces* Genome Database (SGD), 30–40% remained unknown and others only poorly understood. Knowledge acquisition is necessarily a continuum — different experiments are performed at different scales (e.g. high-versus low-throughput) and yield results at different levels of biological detail (e.g. detecting DNA repair versus distinguishing mismatch repair from base excision repair) and confidence (stemming from variation in the quality of assays and the number of replicates performed). For these reasons, the characterization status of gene products does not fall on a simple linear scale. Biologists often make qualitative judgment calls to designate individual gene products as “novel”, “barely characterized”, or “relatively well-characterized”. While this serves the purposes of individual researchers working on a gene-by-gene basis, a more quantitative and objective approach is required to summarize the status of functional characterization for an entire proteome, and to facilitate cross-species comparisons.

### Metrics to describe functional characterization levels

To develop workable metrics for the status of the functional annotation of a given proteome, we have exploited GO annotation [7], and illustrate this scheme using the proteomes of *S. cerevisiae, S. pombe*, and human as examples. Since the functional attributes of gene products of diverse species are routinely described using GO, these metrics are widely applicable.

The GO Molecular Function (MF) ontology describes molecular-level activities of gene products (such as catalytic, transporter, and receptor activities). GO Biological Processes (BP) refers to ordered assemblies of molecular functions representing physiological roles of gene products (e.g. involvement in cytokinesis or DNA replication). GO Cellular Component (CC) provides the locations of gene products (organelles, complexes, etc.). Determining each annotation type relies on different experimental techniques, yielding complementary results and insights. We might know a gene product’s molecular function (MF) but not the physiological context in which that product is involved (BP). As an example, knowing that a protein is a kinase (MF) is not very informative until we find that a deletion mutant of the gene encoding that protein is defective in meiotic nuclear division (BP) and the protein itself localizes to chromatin (CC).

### Annotation coverage for proteins by Gene Ontology aspect

One simple way to quantify the degree to which the function of a given gene product has been characterized is to report annotation to one, two, or all three GO aspects (MF, BP, CC). GO aspect coverage provides a simple metric that is accessible for any species by counting gene products with (or without) annotation to each aspect. Fig 2 shows coverage by ontology aspect for human, fission yeast, and budding yeast proteins. From this viewpoint, we can quickly assess the number of gene products annotated to all three aspects of GO (usually well-characterized), or none at all (uncharacterized), and assess what types are absent for a species.

**Figure 2:**
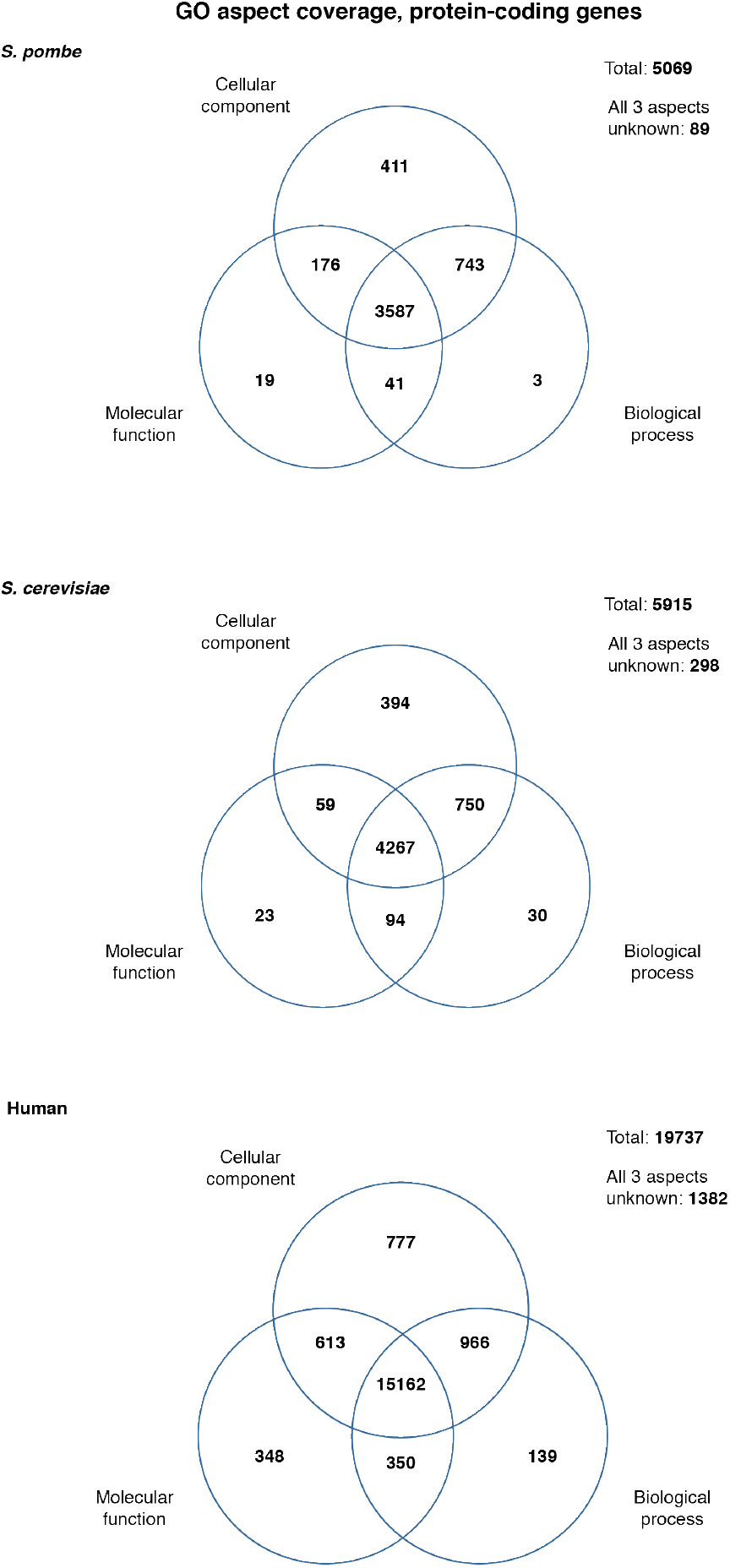
GO aspect coverage of budding yeast, fission yeast, and human proteins. Venn diagrams indicate the number of protein coding gene products annotated to each Gene Ontology aspect (Biological Process, Molecular Function, Cellular Component). Data sources: *S. pombe*, PomBase 25 September 2018; *S. cerevisiae*, YeastMine [28] 25 September 2018; human, HumanMine [29] and GO repository [30], both 26 September 2018.

### Known physiological function: GO slim process coverage

Although informative, the activities captured by MF and the localizations described by CC make only a limited contribution to knowledge about a gene product’s characterization if taken in isolation. In contrast, the physiological context provided by biological process annotation reveals more about the role of a gene product in an organism’s biology and therefore provides a useful benchmark for preliminary characterization.

To use this information as a measure of the progress in a protein’s functional characterization, we use the annotation overviews provided by tailored GO term subsets known as “GO slims” [8]. We created a biological process slim set covering as many annotated gene products as possible, while remaining informative about the physiological context in which they operate. As our starting point, we took the fission yeast GO biological process slim developed at PomBase over 8 years [9], which provides excellent coverage of informative cellular processes for fission yeast (99.4% of annotated proteins with a known process), and minimizes overlap between terms. The PomBase slim aims to demonstrate the distribution of processes within distinct “modules” of biology (cytokinesis, tRNA metabolism, DNA replication etc.) [10, 11], and therefore excludes overly general biological process terms, such as “metabolism” or “cellular component organization”, that would increase coverage at the expense of specific context. Terms that recapitulate activities in the MF ontology (e.g. “protein phosphorylation”) or describe phenotypic observations but do not correspond to a specific physiological role for a gene product (e.g. “response to chemical”) are also excluded. We then extended the 53-term PomBase slim into a generalized process slim of 115 terms to use in cross-species analysis, as summarized in Fig 3A.

**Figure 3:**
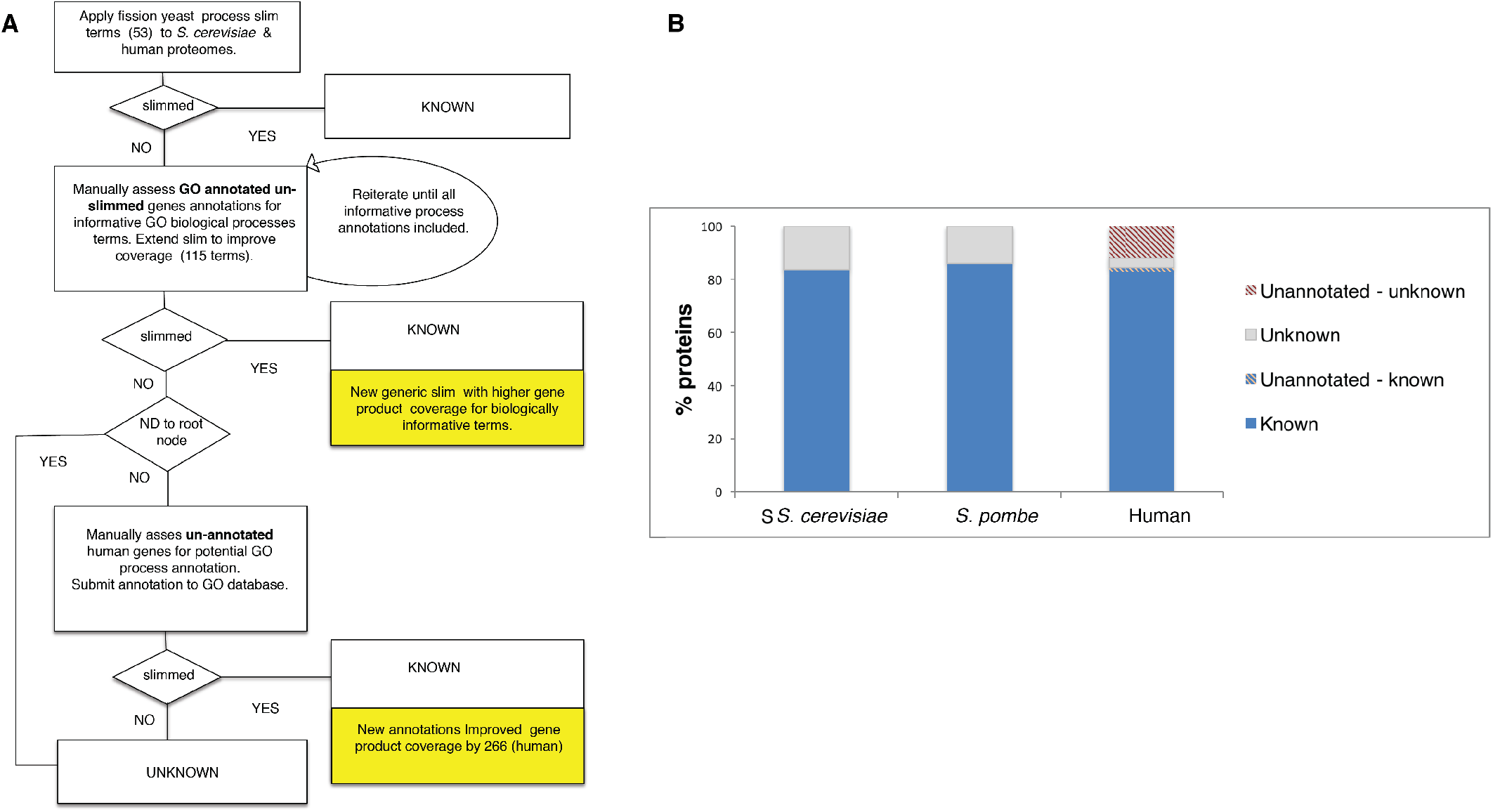
GO slim analysis of budding yeast, fission yeast, and human proteins. **A**: Generic GO biological process slim set creation flowchart. The fission yeast GO biological process slim [9] was applied to human and *S. cerevisiae* protein sets, and then iteratively extended to improve coverage by adding terms. Some processes were swapped (e.g. “cytoplasmic translation” in the fission yeast slim for more general “translation”) to accommodate the less specific annotation available in other species. Overly broad terms (e.g. “metabolism”) and terms representing activities (molecular functions) in the biological process ontology (e.g. “phosphorylation”) were excluded from the generic slim set. At convergence (the point where no additional informative terms could be identified for gene products with biological process annotations), proteins annotated to slim terms were classified as “known” (4393 *S. pombe*; 4936 *S. cerevisiae*; 16354 human). The remaining proteins with uninformative processes were classified as unknown, along with those already identified as unknown by annotation to the root node with evidence code ND (No Data). Manual assessment of the remaining human proteins with no GO biological process annotation added 266 proteins, bringing the “known” total to 16620. Final “unknown” protein totals are 676 in *S. pombe*, 978 in *S. cerevisiae*, and 3117 in human. The set of GO slim terms is available in Supplementary table S1. **B**: Proportions of proteins with known GO slim biological role. For all three species, “known” proteins have annotation to at least one term from the GO slim set (see A), and “unknown” proteins do not. Because the human proteome includes some proteins that lack annotation in the GO database, the proportions of unannotated proteins that we found to be known (i.e. annotatable) and unknown are indicated separately. All protein datasets exclude dubious proteins and transposons. Analysis was performed using GOTermFinder [31], with GO data from 25/09/2018 and the GO slim created as described in A. Input protein lists are available in Supplementary tables S2 (*S. pombe*), S3 (*S. cerevisiae*), and S4 (human). GOTermFinder output is available in Supplementary tables S5 (*S. pombe*), S6 (*S. cerevisiae*), and S7 (human).

For any annotation-based metric, it is important to distinguish *unknown* (or unstudied) from *unannotated* gene products. Here, *unknown* gene products are defined as those that have been evaluated by curators and have no annotation to any BP slim terms (these gene products are annotated to the root term “biological process” with the evidence code “no data (ND)”). *Unannotated* are those not explicitly indicated as unknown but which, nevertheless, have no annotation from experiment or inference. Because all fission yeast and budding yeast genes have been systematically assessed using all available data, any gene products lacking specific GO annotations can confidently be deemed to have unknown biological roles. For the human proteome, manual inspection of the unannotated proteins revealed that many can actually be annotated to a biological process based on experimental data in the literature or by homology-based inference, and thus classified as characterized. To make this knowledge available, we manually curated 1235 GO annotations for 598 human proteins from 348 publications, including biological process assignments for 266 previously unannotated proteins. These annotations will be submitted to the GO Consortium for inclusion in the human GO annotation dataset. Fig 3B shows the proportions of the *S. pombe, S. cerevisiae*, and human proteomes that are *known* (i.e. annotated to informative biological process terms), *unannotated, annotable*, or *unknown*. See Supplementary Data for GO slim term IDs (Table S1), input protein lists (Tables S2–S4), GO slim outputs (Tables S5–S7), and unknown gene lists (Tables S8–S10).

## Why do these proteins remain unstudied?

Our GO slim-based characterization metric confirms the impression from simpler metrics that, for the two model yeasts and human, about 20% of proteins lack physiologically informative descriptions. Why do so many proteins, many of them conserved, remain unstudied? Below, we consider biological and sociological/cultural factors that contribute to the apparent lack of interest in these unknown proteins.

### Biological bias

One factor influencing gene characterization is simply how easily one gene’s contribution to an organism can be detected. Deletion mutants of essential genes have a clear phenotype that indicates an important function — for yeasts, the failure to grow on rich media. As a consequence, these genes, and the core processes in which they participate, are well characterized. For example, only 24 of the genes in the fission yeast unknown set are essential in rich media. Changes in cell morphology are also readily identifiable phenotypes. Visual screening and analysis of the fission yeast genome-wide deletion collection for morphology phenotypes under standard laboratory conditions found obvious abnormalities for only 10% of 643 genes of unknown function [12]. The most commonly used experimental conditions, designed as they are to maximize cell growth, can hide environment-dependent roles. Many more of the 676 currently uncharacterized (per Figure 3B) fission yeast genes are associated with growth or viability phenotypes upon specific chemical challenges (26.1%) than under standard laboratory conditions (3.6%) (PomBase [10, 11], queries run 11/11/2018). In budding yeast, only 34% of all deletion mutants display a growth phenotype under standard laboratory conditions, whereas 97% of all genes are essential for optimal growth in at least one condition when assayed under multiple chemical or environmental perturbations [13].

Protein characterization has traditionally emphasized core biological processes over those that reflect interactions with the environment. However, analysis of the proteins that have recently (2016–2018) been removed from the fission yeast “conserved unknown” set because their functions have been determined reveals that they most often participate in environment-responsive processes such as signalling, detoxification, proteostasis, lipostasis and mitochondrial organization (Table 1). Many of these functions are associated with the age-related accumulation of damaged or misfolded proteins, which become debilitating over time. In humans, such functions are implicated in neurodegenerative diseases, such as Alzheimer’s and motor neuron diseases [14], which underscores the importance of making strenuous efforts to elucidate the functions of the remaining “conserved unknowns”. It should also be pointed out that the ecology of yeasts is poorly understood, and we postulate that many unknown proteins function in aspects of life that are not normally probed in the laboratory (e.g., interactions with pathogens, or survival within insect vectors). We anticipate that a greater variety of experimental conditions will supply more information about the unknown gene products catalogued here to reveal the roles of many gene products in processes fundamental for human health.

**Table 1:** GO slim classification of conserved *S. pombe* proteins characterized between 2016 and 2018.

| Process | Proteins |
| --- | --- |
| **Membrane biology** |  |
| lipid metabolism | 15 |
| transmembrane transport | 9 |
| vesicle-mediated transport | 9 |
| organelle localization by membrane tethering | 4 |
| other membrane organization | 3 |
| other ER processes | 2 |
|  | 42 |
| **Communication** |  |
| signaling | 9 |
| transcription | 6 |
| chromatin organization | 1 |
|  | 16 |
| **Catabolism and detoxification** |  |
| detoxification | 25 |
| protein catabolism | 5 |
| apoptotic process | 4 |
| DNA repair | 3 |
| nucleobase-containing compound catabolism | 5 |
| autophagy | 1 |
| mannose catabolism | 1 |
|  | 44 |
| **Mitochondrial processes and energy** |  |
| mitochondrial gene expression | 4 |
| mitochondrial organization | 3 |
| energy generation | 4 |
|  | 11 |
| **Other processes** |  |
| tRNA metabolism (cytosolic) | 3 |
| ribosome biogenesis (cytosolic) | 5 |
| mRNA metabolism | 2 |
| cytoplasmic translation | 2 |
| cytoskeleton organization | 3 |
| protein folding (cytosolic) | 1 |
| protein complex assembly | 1 |
| nucleocytoplasmic transport | 1 |
| chromosome segregation | 1 |
| amino acid metabolism | 2 |
| cofactor metabolism | 1 |
| other | 5 |
|  | 27 |
| **TOTAL** | 140 |

### Research bias

In fission yeast, the majority of new knowledge over the past decade provides increasing detail for previously described proteins. This bias towards studying already-known proteins is not peculiar to yeast research. In their essay “Too many roads not taken” [15], Edwards *et al*. observe that 70% of human protein research still focuses on the 10% of proteins known before the human genome was sequenced. Although few studies have explored the causes of the observed emphasis on known proteins, we can identify a number of plausible contributing factors, which are largely borne out by a recent large-scale analysis of publications on human genes [16]. First, the complexity of biology demands that investigators narrow their study targets to a manageable range. Researchers with established interests in specific topics thus naturally focus their work where they have deep knowledge, and extend their studies to novel genes only if a strong lead emerges, for instance, from work in another species or from a data-mining approach that implicates them in biological processes already under investigation. Indeed, Stoeger *et al*. find that research in model organisms strongly influences the initial study of individual human genes. Both papers also highlight pragmatic considerations, notably the availability of research tools, and socio-political factors including career timelines, funding priorities, and peer review, all of which exacerbate the tendency to avoid the wholly unknown. Risk-averse funders and reviewers tend not to favor long-range strategies aimed at genes without an existing functional context for fear of diverting resources towards targets whose significance is not guaranteed. Without a shift in perspective, proteins without any existing functional annotation will continue to be neglected, to the detriment of basic and applied biomedical research. Stoeger *et al*. note that current research is not only slow to cover novel genes, but also “can significantly deviate from the actual biological importance of individual genes”.

## Classifying the conserved unknown proteins, or: What lies undiscovered?

The stubborn core of remaining proteins of unknown function are often dismissed as species-specific, but we have often found otherwise, and we can no longer afford to sweep these proteins under the carpet. Therefore, to provide further insight into why some gene products elude physiological characterization, we present a case study using the set of 410 fission yeast proteins of unknown physiological role that have orthologs outside the *Schizosaccharomyces* clade (the “conserved unknowns”). We classify these 410 proteins according to a range of orthogonal biological attributes (including taxonomic conservation, identification of a catalytic fold or domain, cellular localization, viability). Fig 4 presents the subset of 200 proteins in this group which are conserved outside fungi (in vertebrates, archaea, or bacteria; details for the full set of 410 proteins are provided in Supplementary Figure S1 and Table S11). The number of conserved unknown proteins that play an essential role under permissive growth conditions is disproportionally low (4.4%; 18/410), and almost half of these 18 essential proteins are localized to the mitochondria or the endoplasmic reticulum. Only 8 essential genes are conserved in vertebrates (all are organellar), and only a single protein out of 53 conserved between yeast and prokaryotes (but not vertebrates) is essential. A substantial proportion of the 200 proteins conserved outside the fungi (76) are absent from *Saccharomyces* due to the well-documented lineage-specific gene losses in its evolutionary history [17]; characterized gene products with this taxonomic distribution are most highly enriched for chromatin organization and mRNA metabolism. Unknown mitochondrial and endomembrane system proteins are enriched for proteins with transmembrane domains (59/114). Unknown nuclear proteins are predicted to include more transferases than other enzymatic activities; the set of nuclear proteins shared only with eukaryotes includes 19 non-catalytic domains of unknown function (DUFs) and 12 protein-protein interaction domains (e.g. WD, ankyrin, or TPR) that frequently function as scaffolds for protein complexes [18]. This multi-factorial classification can support the prediction of likely physiological roles that can be experimentally tested.

**Figure 4:**
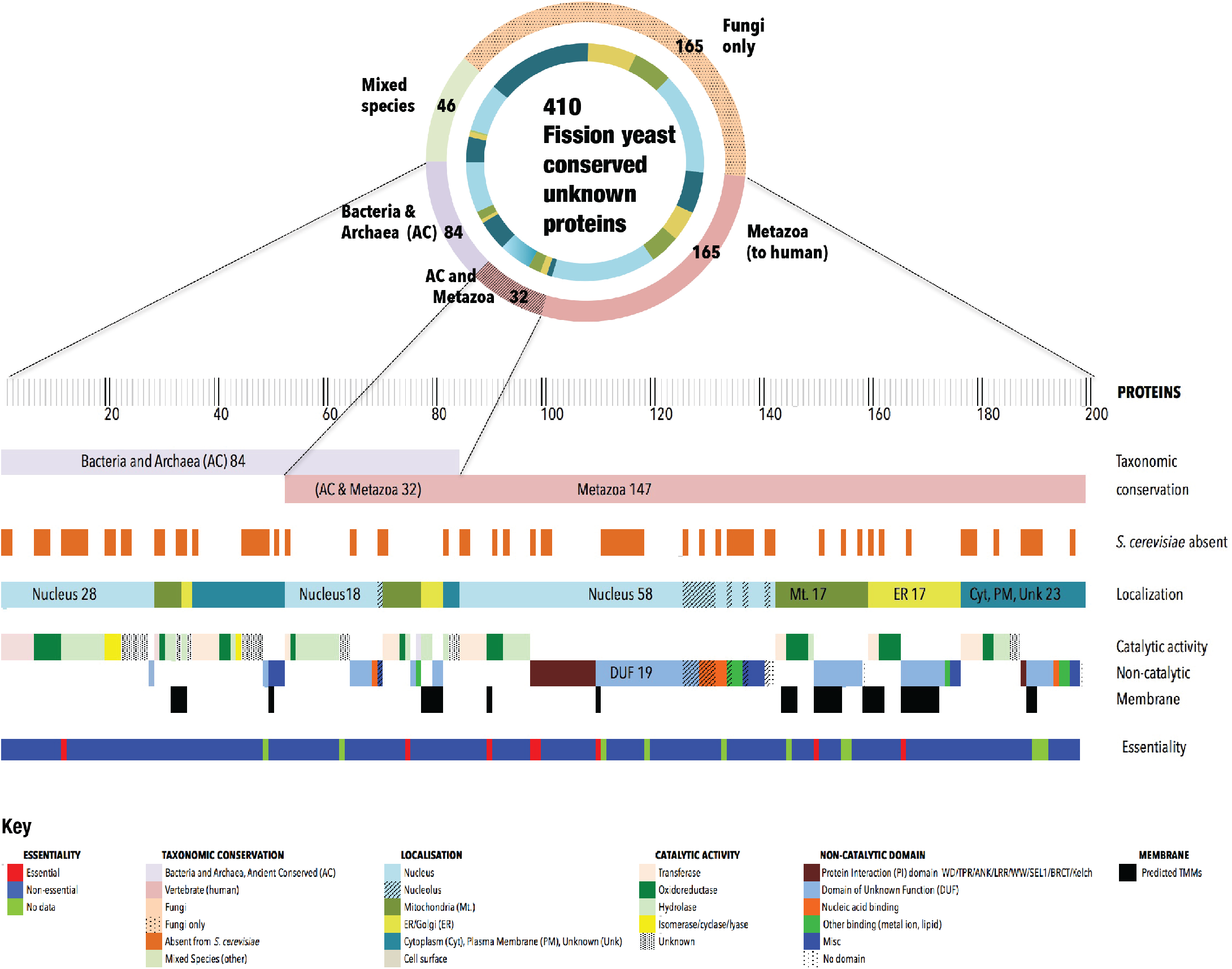
Taxonomic conservation and features of unknown proteins. Classification of 210 conserved unknown fission yeast proteins along various axes. PomBase curators manually assign protein-coding genes to one of a set of broad taxonomic classifiers [11, 32]. PomBase also maintains manually curated lists of orthologs between *S. pombe* and *S. cerevisiae*, and between *S. pombe* and human, three eukaryotic species separated by ca. 500–1000 million years of evolution. In combination, these inventories can be used to identify conservation across taxonomic space at different levels of specificity. Of the fission yeast “unknown” protein-coding genes, 410 are conserved outside the *Schizosaœharomyœs* clade. Of these, 210 are present either in fungi and vertebrates, or in fungi and prokaryotes (data from PomBase manual assignments, queried on 31/07/2018). Proteins were classed as catalytic or non-catalytic based on protein domain, fold, clan or superfamily membership, using InterPro [18] and GO [7] assignments. Cellular locations using GO annotation are available for most of the unknown proteome based on a genome-wide localization study and inference from other models [19]. Viability data come from large-scale screens reported by Kim *et al*. [20] and Chen *et al*. [21]. The fission yeast “conserved unknown” protein set [33] is reviewed continually for new functional data.

## Strategies to link unknown genes to broad cellular roles

Despite almost a century of gene product-specific biochemical and genetic interrogation, and two decades of post-genomic research, a large number of proteins conserved from yeast to human still have no known biological role. Broadly conserved unknown eukaryotic proteins can be assumed to have important cellular roles conserved over 500 million years of evolution. It is, therefore, remarkable that this gene set has hardly reduced over the past decade. In this period, familiar genes have been studied in ever-greater depth, presumably at the expense of the characterization of genes of hitherto unknown function (e.g. over 33,000 papers with “p53” in title have been published since 2007). Assuming a diminishing return for studies on highly characterized proteins, investigations on unstudied proteins will have a relatively higher impact that is likely to outweigh the considerable initial efforts required to place them within the context of current knowledge.

To jump-start renewed progress in unknown gene characterization, two major stumbling blocks must be overcome: One is to identify the cohort of genes of unknown function, and the other is to develop mechanisms to bring the proteins that they encode to the attention of interested researchers. Here, we provide a framework that uses a generally accessible set of criteria based on manually curated data to identify and classify unstudied proteins, which could easily be extended with additional criteria for further annotation specificity in future iterations. The construction of inventories of unknown proteins will ultimately depend on accurate and complete functional annotation of all genes of the major model species.

Commentators on genome-scale research have long recognized that, in order to fully describe an organism’s protein complement, it will be necessary to deploy parallel experimental and computational methods, at both large and small scales [4, 6]. Understanding how investigatory biases and the characteristics of particular gene sets have converged to prevent characterization will help us to identify the most promising routes to uncovering unknown functions. These, and other factors that contribute to the neglect of the characterization of conserved gene products that are likely to have novel biological roles, deserve further in-depth consideration. It is likely that, to fill the persistent knowledge gaps represented by the roughly 20% of proteins that remain uncharacterized, a creative combination of existing and emerging experimental and *in silico* methods will be required, as well as an increased awareness among the scientific community of the value of a full proteome description. Because basic knowledge at the cellular level provides the building blocks of translational research, drug discovery, personalized medicine, metabolomics, and systems biology, comprehensive proteome characterization underpins the success of numerous and diverse endeavors across all of the biological and medical sciences.

## Supplementary materials

- Figure S1. Taxonomic conservation and features of unknown proteins. Classification of 410 conserved unknown fission yeast proteins (PomBase manual taxonomic conservation assignments, queried on 31/07/2018) along various axes. Proteins were classed as catalytic or non-catalytic based on protein domain, fold, clan or superfamily membership, using InterPro [18] and GO [7] assignments. Cellular locations using GO annotation are available for most of the unknown proteome based on a genome-wide localization study and inference from other models [19]. Viability data come from large-scale screens reported by Kim *et al*. [20] and Chen *et al*. [21].
- Table S1. GO IDs for biological process GO slim set used in Figure 4 analysis.
- Table S2. *S. pombe* proteins used in Figure 4 analysis (PomBase systematic IDs).
- Table S3. *S. cerevisiae* proteins used in Figure 4 analysis (SGD ORF names and UniProtKB accessions).
- Table S4. Human proteins used in Figure 4 analysis (UniProtKB accessions).
- Table S5. *S. pombe* GO slim results from Figure 4 analysis.
- Table S6. *S. cerevisiae* GO slim results from Figure 4 analysis.
- Table S7. Human GO slim results from Figure 4 analysis.
- Table S8. Unknown *S. pombe* proteins from Figure 4 analysis.
- Table S9. Unknown *S. cerevisiae* proteins from Figure 4 analysis.
- Table S10. Unknown and unannotated human proteins from Figure 4 analysis. A, proteins that had no GO biological process annotation when analyzed, but can be annotated based on available literature. B, proteins remaining unknown following analysis (annotated to the GO biological process root, or lacking both annotation and data that would support annotation).
- Table S11. List of genes classified in Figure S1.

## Acknowledgements

Funding: This work was supported by the Wellcome Trust (grant 104967/Z/14/Z to SGO). Author contributions: VW conceived the project and wrote the initial draft. VW, AL, and MAH prepared figures. All authors contributed to the discussion of ideas and manuscript revisions, and read and approved the final manuscript. Competing interests: The authors declare no competing interests.

